# PRADA-DTI: A Prototype-Retrieval Augmented Domain-Adaptation Framework for Drug-Target Interaction Prediction

**DOI:** 10.64898/2026.01.05.697600

**Authors:** Jie Zhu, Tianxu Lv, Xiang Pan

## Abstract

Drug target interaction (DTl) prediction is a fundamental task in computational drug discovery. However, most existing DTI models assume a static learning environment, whereas real-world biomedical data are dynamic, characterized by the continuous emergence of new protein families and interaction patterns. This poses major challenges for model generalization and continual adaptation, especially under privacy and data-access constraints. To address these issues, we introduce PRADA-DTI, a retrieval-augmented, parameter-efficient framework for domain-incremental DTI learning. Built on a shared physicochemical backbone, PRADA-DTI learns domain-specialized prompts for representation-level modulation, coupled with Low-Rank Adaptation (LoRA) modules for parameter-space adaptation. At inference, protein embeddings query a compact prototype memory without storing raw molecular data, retrieving similar domains to dynamically compose the relevant prompts and LoRA parameters without requiring domain labels. This retrieval-guided composition enables continual learning from new protein domains while mitigating catastrophic forgetting on previous ones. On BindingDB and BIOSNAP benchmarks, PRADA-DTI substantially outperforms state-of-the-art continual learning and parameter-efficient baselines in both predictive accuracy and forgetting mitigation with minimal parameter updates. Interpretability analysis through residue-level attribution visualization demonstrates that the model correctly attends to binding pocket regions across different protein domains, confirming that the retrieval-guided adaptation mechanism captures biologically relevant structural patterns. These results demonstrate the effectiveness of retrieval-augmented parameter adaptation for continual drug discovery.

## 1 Introduction

Drug–target interaction (DTI) prediction plays a central role in computational pharmacology, enabling large-scale virtual screening through in-silico modeling of drug compounds and protein targets [1, 2]. Early approaches relied on chemical or sequence similarity and matrix factorization [3, 4], while more recent models utilize deep learning to encode heterogeneous molecular inputs. Sequencebased models such as DeepDTA and MolTrans [5, 6] employ CNNs or Transformers on SMILES strings and amino acid sequences, whereas structure-based methods like GraphDTA process molecular graphs [7]. Advanced variants further incorporate 3D conformations or pocket-level features [8, 9]. However, most DTI models assume static independent and identically distributed data, where drug-protein pairs are sampled from a shared distribution. In real-world settings, distribution shifts across species or protein families are common, and prior studies on benchmarks such as TDC have shown that such shifts significantly impair generalization, causing performance degradation when test proteins come from dissimilar domains [2, 10, 11].

To address these challenges, we formulate DTI prediction as a domain incremental learning problem [12, 13], where protein domains appear sequentially and the model must update without replay or prior exposure. This setting aligns with real biochemical pipelines in which new protein families are routinely introduced. Although continual learning (CL) has been investigated extensively in vision and NLP [14, 15, 16], its application to molecular prediction is still relatively underexplored. Drug molecules and proteins exhibit complex structural organization, including multi-scale graph topology and long-range spatial dependence, which can challenge continual adaptation and limit the feasibility of reusable memory mechanisms. These considerations motivate the need for lightweight and modular forms of adaptation. Parameter-efficient fine-tuning (PEFT) methods such as LoRA [19] and prompt tuning [20, 21] offer compact domain adjustments while keeping the backbone fixed; however, protein-ligand interactions are often sensitive to subtle conformational variation, and PEFT may show reduced robustness when generalizing across proteins with substantial structural differences. Moreover, standard PEFT techniques do not typically include mechanisms for retrieving or reusing previously learned domain information during inference. Inspired by retrieval-augmented frameworks such as RAG [22], we seek an approach that incorporates retrieval signals into domain-adaptive tuning.

**Fig. 1:**
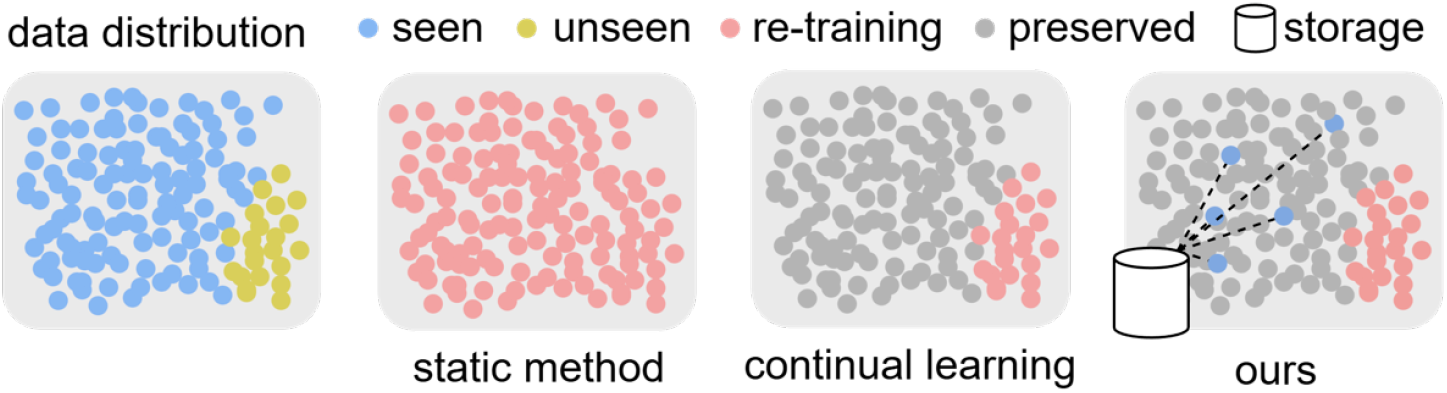
Comparison of static training, conventional continual learning, and our prototype-guided approach from the data perspective.

Building on this idea, we develop PRADA-DTI, a continual learning framework that couples prototype-based retrieval with parameter-efficient adaptation. The model keeps a shared physicochemical graph backbone fixed and attaches lightweight LoRA and prompt modules to capture domain-specific adjustments. After training on each domain, representative protein prototypes are stored in a compact memory. During inference, the query protein retrieves its nearest prototypes, and the associated LoRA and prompt parameters are combined through similarity-weighted aggregation. This retrieval-guided composition enables the model to select domain-relevant adaptations without task identifiers or data replay, providing a flexible and privacy-preserving mechanism for handling unseen protein domains and supporting continual generalization across evolving biochemical distributions.

### PRADA-DTI provides the following contributions

- **Retrieval-augmented continual adaptation**. We introduce a retrieval-driven mechanism that selects domain-relevant prototypes for each protein query, enabling task-ID–free and replay-free adaptation.
- **Dual-level parameter-efficient tuning**. PRADA-DTI combines prompt tuning at the feature level with LoRA at the parameter level, offering complementary domain-specific adjustments across evolving protein domains.
- **Modular and scalable architecture**. The physicochemical graph backbone remains fixed, and fewer than 3% of parameters are updated per domain, allowing the framework to scale to many domains.
- **Consistent empirical improvements**. PRADA-DTI achieves higher accuracy and lower forgetting than continual learning and PEFT baselines on BIOSNAP and BindingDB.

**Fig. 2:**
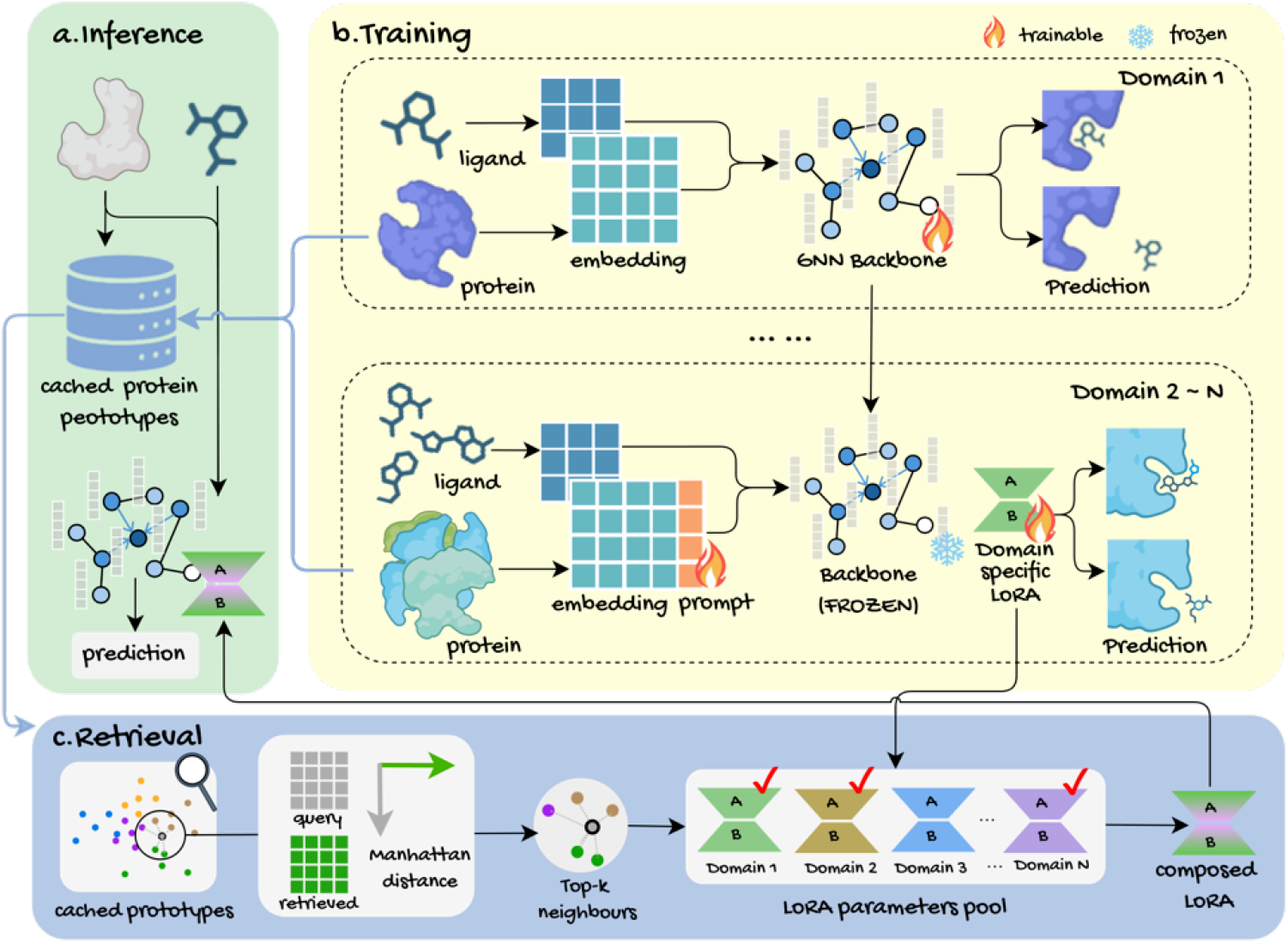
Overall pipeline of our framework. (a) Inference. Protein embeddings query a prototype memory to retrieve similar domains. Retrieved prototypes guide the composition of LoRA parameters for domain-aware prediction.(b) Training. The GNN encoder is first trained and then frozen. For each new domain, only LoRA and prompt modules are updated for lightweight domain-specific adaptation.(c) Retrieval. Protein queries match cached prototypes via Manhattan distance. Top-k nearest neighbors select corresponding LoRA modules, which are composed at test time.

## 2 Methodology

We propose a retrieval-augmented parameter adaptation framework for domain-incremental DTI prediction. The method combines prototype-based retrieval, domain-specific prompts, and dynamically composed LoRA adapters, while employing a lightweight shared prediction head that avoids explicit task identifiers.

### 2.1 Problem Definition and Notation

We study domain-incremental DTI prediction, where a model observes a sequence of domains 𝒟_1_, …, 𝒟_*T*_ and is trained on each domain 𝒟_*t*_ without access to data from earlier domains. Each domain consists of samples (*G*_*d*_, *G*_*p*_, *y*), where *G*_*d*_ and *G*_*p*_ denote drug and protein graphs, and *y* ∈{0, 1}indicates interaction. Domains are disjoint in protein space, reflecting realistic distribution shifts across protein families and resulting in a non-IID learning setup.

The model follows standard domain-incremental constraints: *no rehearsal buffer, no domain identity* during inference, and *single-pass* updates per domain. The learning objective is to maximize the average performance over all domains:

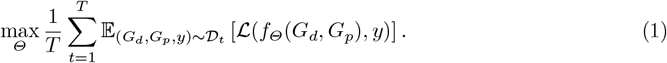

This setting naturally leads to catastrophic forgetting, as updates on later domains may interfere with previously learned protein-level knowledge.

### 2.2 Frozen Molecular Interaction Backbone

To maintain stable representations across domains, we freeze a shared molecular interaction backbone that encodes drug-protein pairs using graph-based models.

#### Protein encoder

Each protein is represented with residue-level and atom-level graphs. Residue embeddings from ESM-2 [23] are enriched with local atomic graph features to produce a hierarchical protein embedding *E*_*p*_ ∈ℝ^*d*^.

#### Drug encoder

Drug molecules are encoded as atom–group graphs. A pretrained ChemBERT model [24] generates contextual node features, which are aggregated by a GNN to obtain a molecular embedding *E*_*d*_ ∈ ℝ^*d*^.

#### Cross-modal interaction

Drug and protein embeddings are fused using a multi-head cross-attention module:

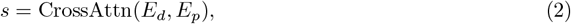

yielding an interaction score *s* ∈ ℝ.

All backbone parameters remain frozen during continual training, and domain-specific adaptation is handled by lightweight external modules introduced in the following sections.

### 2.3 Low-Rank Domain-Specific Adaptation

Low-rank adapters provide a parameter-efficient way to incorporate domain-specific updates while keeping the backbone frozen. Given a pretrained weight matrix *W* ∈ ℝ^*d*×*d*^, LoRA models the update as:

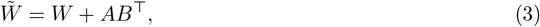

where *A, B* ∈ ℝ^*d*×*r*^ are trainable low-rank matrices with *r* ≪*d*. Only *A* and *B* are optimized, enabling lightweight gradient updates without modifying shared parameters.

We insert LoRA modules into selected layers of the classification head, protein encoder, and interaction modules. The insertion points are determined through a sensitivity analysis of parameter changes across domains (details in **Appendix C**).

Each domain 𝒟_*t*_ maintains its own LoRA parameters {*A*_*t*_, *B*_*t*_}, providing isolated parameter-space adaptations, reducing interference in rehearsal-free continual learning.

### 2.4 Prompt-Based Feature Adaptation

Prompt tuning provides complementary feature-level adaptation to LoRA. For each domain 𝒟_*t*_, we introduce a trainable prompt matrix *p*_*t*_ ∈ ℝ^*l*×*d*^ and prepend it to the ESM2-derived residue embeddings of the protein encoder:

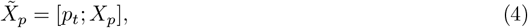

where *X*_*p*_ denotes the original ESM2 feature sequence. These prompts act as soft prefix tokens that condition the protein encoding process from the input layer onward.

During training on 𝒟_*t*_, the prompt vectors are optimized jointly with the corresponding LoRA parameters. Prompt tuning adjusts feature-level representations, while LoRA modifies deeper transformation layers, providing complementary and lightweight domain-specific adaptation.

### 2.5 Prototype-Retrieval for Dynamic Parameter Adaptation

To enable domain-aware adaptation without task identifiers or rehearsal data, we introduce a prototype-retrieval mechanism that selects relevant domain knowledge at test time and uses it to compose lightweight adaptation parameters. Instead of combining prototypes directly, similarity to stored prototypes determines how LoRA and prompt modules from previously seen domains are merged for the current query.

#### Prototype Construction

After training on each domain 𝒟_*t*_, protein embeddings from the frozen ESM-2 encoder are clustered using *k*-means to obtain prototype vectors 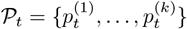. These prototypes summarize the structural distribution of proteins in 𝒟_*t*_ and are stored in a global memory ℳ_*P*_ =∪_*t*_. 𝒫_*t*_ Because they are derived from high-level embeddings rather than raw sequences, the prototypes serve as anonymized and task-agnostic representations.

#### Prototype Retrieval

Given a query protein embedding *E*_*p*_, Manhattan distance is computed between *E*_*p*_ and each stored prototype. The similarity score for prototype *p*^(*i*)^ is defined as the negative distance:

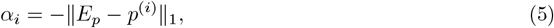

so that closer prototypes receive larger similarity values. A *K*-nearest neighbor search retrieves the top-*K* most similar prototypes 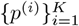 and their similarity scores are normalized by:

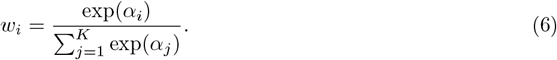

Each prototype corresponds to its originating domain 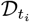 and is linked with its LoRA parameters 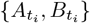 and prompt vector 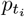.

#### Parameter Composition

The adaptation parameters for the query protein are obtained through similarity-weighted mixing:

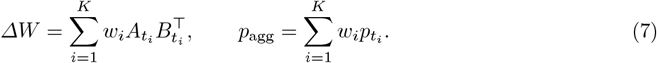

This retrieval-guided composition interpolates among relevant domains in parameter space, producing adaptation parameters tailored to the query protein rather than switching between discrete tasks.

The prototype-retrieval mechanism thus turns domain adaptation into a continuous, similarity-weighted parameter selection process. It enables soft, data-free adaptation to unseen or mixed-domain samples, mitigates forgetting, and remains compatible with the frozen backbone and parameter-efficient modules.

## 3 Results

### 3.1 Experimental Setup

#### Datasets

We evaluate PRADA-DTI on two widely used DTI benchmarks, BIOSNAP and BindingDB, under a domain-incremental setting where protein domains are introduced sequentially, without access to task identifiers or rehearsal buffers. BIOSNAP contains experimentally validated binary drug-target interactions curated from diverse biological databases, while BindingDB provides largescale affinity measurements based on experimentally determined binding activities.

To simulate domain-incremental scenarios, proteins in each dataset are clustered into multiple domains using the Leiden community detection algorithm applied to their ESM2 embeddings. We conducted a grid search over the resolution and neighborhood size (*k*) of the k-nearest-neighbor graph and adopted resolution = 0.001 and *k* = 60, yielding 15 and 25 domains for BIOSNAP and BindingDB respectively. These domains are non-overlapping and well-separated in the embedding space. Full details on the clustering process are provided in Appendix A.

#### Baselines and Evaluation Metrics

We compare PRADA-DTI against several representative baselines: **EWC** [26], a regularization-based continual learning method that constrains changes to important parameters to reduce forgetting; **LwF** [27], a knowledge distillation approach that encourages the model to retain previous knowledge by matching current outputs to earlier outputs when learning new domains; **TPP** [29], a graph continual learning method that leverages Laplacian smoothing-based task prototypes for accurate task identification and introduces graph prompting to mitigate forgetting without data replay; **HPNs** [30], a hierarchical prototype network that extracts multilevel prototypes to represent expanding graphs, enabling adaptive feature selection and refinement for new categories while effectively preventing forgetting; and **MSCGL** [31], a continual graph learning method that combines neural architecture search and group sparse regularization to adapt to evolving graph structures and mitigate forgetting across sequential domains. All models utilize the same PSICHIC backbone and are trained under consistent settings to ensure fair comparison.

For evaluation, we report four metrics for all models: accuracy (ACC), area under the ROC curve (AUC), average forgetting (AFGT, measuring the performance drop on previous domains), and backward transfer (BWT, quantifying the influence of new domain learning on past knowledge).

#### Implementation Details

Each input sample consists of a drug SMILES string and a protein amino acid sequence. Drugs are processed into molecular graphs using RDKit, and then embedded using a pre-trained ChemBERT encoder. Proteins are embedded using the ESM2 model. All methods share the same PyTorch-based PSICHIC backbone, which remains frozen during continual learning. Adaptation is achieved via LoRA and prompt-based modules. Models are trained with binary crossentropy loss using the AdamW optimizer (initial learning rate 1 × 10^−5^), a 5% warmup, and cosine decay schedule. Each domain is trained for up to 30 epochs, with early stopping based on validation AUC.

During inference, prototype retrieval is performed using Manhattan distance to select the top-3 nearest prototypes with a temperature parameter *τ* = 0.07. All experiments are conducted on a single NVIDIA RTX 4090 GPU (24 GB). Details on prototype retrieval and storage are provided in Appendix A.

#### Training and Inference Protocol

We outline here the training and inference procedure used for PRADA-DTI in the domain-incremental setting.

##### Training Phase

During training on each domain 𝒟;_*t*_, the backbone encoders and fusion module remain frozen. Only the domain-specific LoRA parameters {*A*_*t*_, *B*_*t*_} and the prompt vector *p*_*t*_ are updated using binary cross-entropy loss on DTI labels. The retrieval mechanism is disabled during training to avoid cross-domain information leakage. After completing training on 𝒟_*t*_, protein embeddings from the frozen ESM-2 encoder are clustered to form domain prototypes, which are then added to the global prototype memory ℳ_*P*_.

##### Inference Phase

At inference time, the prototype-retrieval mechanism is activated. Given a test protein, its embedding is computed and compared against all stored prototypes via Manhattan distance. The top-*K* retrieved prototypes determine similarity weights that guide the composition of LoRA updates and prompt vectors into query-specific adaptation parameters. These dynamically assembled modules are subsequently used to compute the final interaction score for the drug–protein pair.

This protocol allows each domain to be learned independently while enabling domain-aware adaptation at test time through retrieval-guided parameter composition. Full implementation details, including optimization schedules, batch sizes, and hardware configurations, are provided in Appendix A.

### 3.2 Main Results

#### Overall performance

As shown in Table 1, PRADA-DTI achieves the highest average accuracy and the lowest forgetting across domains on both datasets. Regularization-based methods restrict parameter updates, and topology-aware graph methods preserve structural consistency, yet both remain limited because continual optimization of shared backbone parameters leads to interference. PRADA-DTI avoids this issue by freezing the backbone and adapting only through lightweight LoRA adapters and prompt vectors. This modular design substantially reduces cross-domain interference and results in negligible forgetting.

**Table 1:** Evaluation of continual learning methods on BIOSNAP and BindingDB under domainincremental settings. Reported metrics include accuracy (ACC), area under the ROC curve (AU-ROC), backward transfer (BWT), and average forgetting (AFGT), where ↑ / ↓ indicate whether higher or lower values are better. Joint denotes a non-incremental upper bound trained on all domains simultaneously and is not included in the final comparison with other methods. For each metric, the best result among all continual learning methods is highlighted **in bold**, and the second-best is underlined.

| Method | Biosnap |  |  |  | BindingDB |  |  |  |
| --- | --- | --- | --- | --- | --- | --- | --- | --- |
|  | ACC(↑) | AUROC(↑) | BWT(↑) | AFGT(↓) | ACC(↑) | AUROC(↑) | BWT(↑) | AFGT(↓) |
| Finetune | 63.86 | 0.5745 | -24.4452 | 23.37 | 55.48 | 0.5379 | -28.4128 | 28.33 |
| Joint | 86.37 | 0.8962 | - | - | 88.45 | 0.9007 | - | - |
| EWC | 66.60 | 0.6278 | -5.2590 | 15.21 | 55.13 | 0.5630 | -12.6739 | 15.84 |
| LwF | 68.86 | 0.6557 | -4.6276 | 14.83 | 58.83 | 0.5879 | -2.6915 | 15.27 |
| MAS | 67.73 | 0.6389 | -11.8623 | 15.02 | 59.16 | 0.5746 | -28.6440 | 14.96 |
| ER | 70.58 | 0.6279 | -19.4684 | 13.51 | 58.83 | 0.5731 | -23.8620 | 24.46 |
| TPP | 71.25 | 0.6732 | -2.0821 | <u>2.46</u> | 65.78 | 0.6028 | -5.5182 | 8.73 |
| MSCGL | <u>76.43</u> | 0.6805 | <u>-0.7914</u> | 2.485 | <u>68.82</u> | 0.6332 | <u>-2.4145</u> | <u>3.76</u> |
| HPNs | 72.23 | <u>0.6932</u> | -3.4895 | 6.82 | 65.47 | <u>0.6478</u> | -5.8971 | 8.41 |
| **Ours** | **82.13** | **0.7842** | **-0.1077** | **0.17** | **70.38** | **0.6915** | **-0.1805** | **0.79** |

#### Forgetting analysis

Figure 3 illustrates that PRADA-DTI maintains an almost flat forgetting curve across all domains. Baseline methods-regardless of whether they are regularization-based or graphstructured-exhibit steadily increasing forgetting, especially under larger distribution shifts. PRADA-DTI stays within a few percentage points throughout, indicating far better stability and long-term retention.

**Fig. 3:**
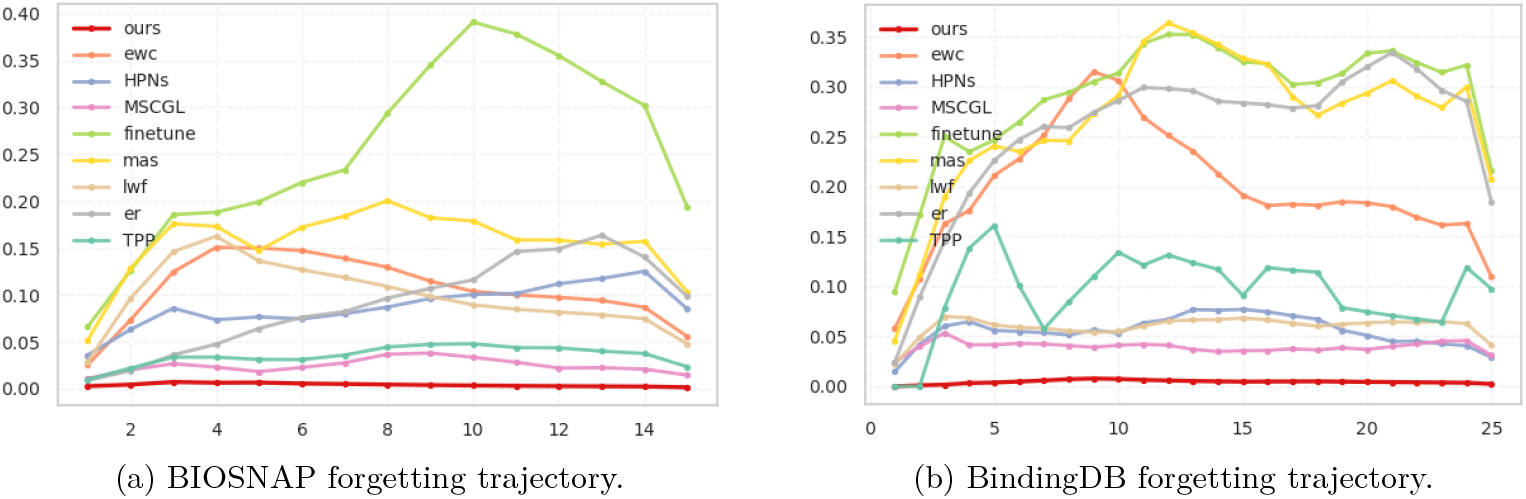
Forgetting trajectories of continual learning methods on BIOSNAP (left) and BindingDB (right). We report relative average forgetting (%) as a function of the number of sequentially learned domains. Each point reflects the average forgetting measured up to that domain under a domain-incremental setting without data replay or task identifiers. Lower curves indicate better forgetting resistance.

### 3.3 Ablation Studies

To understand the effect of each component, we ablate retrieval (R), multi-domain LoRA composition (M), and prompt tuning (P). The results in Table 2 show a clear hierarchy of contributions. Using prompts alone yields only modest improvements, indicating that input-level conditioning is insufficient for capturing domain variation in protein-ligand interactions. Adding the retrieval module leads to a substantial boost in both accuracy and forgetting metrics on both datasets, confirming that prototype-guided parameter selection is the primary driver of effective domain adaptation. Introducing multi-domain LoRA composition further improves performance, showing that interpolating across domain-specific adapters provides a more flexible and expressive parameter space.

**Table 2:** Ablation study of PRADA-DTI on BIOSNAP and BindingDB. We examine the contributions of retrieval (R), multi-LoRA (M), and prompt tuning (P) by selectively enabling each component.” √”.indicates that the corresponding module is used; “×” denotes ablation. Metrics include accuracy (ACC), Area Under the ROC Curve (AUC), backward transfer (BWT), and average forgetting (AFGT).

| Biosnap |  |  |  |  |  |  | BindingDB |  |  |  |
| --- | --- | --- | --- | --- | --- | --- | --- | --- | --- | --- |
| R | M | P | ACC | AUC | BWT | AFGT | ACC | AUC | BWT | AFGT |
| × | × | ✓ | 63.49 | 57.57 | -22.86 | 20.46 | 58.68 | 56.67 | -28.75 | 24.84 |
| × | ✓ | × | 65.56 | 63.33 | -16.55 | 15.56 | 60.25 | 58.52 | -10.23 | 15.96 |
| ✓ | × | × | 75.43 | 67.66 | -6.08 | 5.16 | 67.73 | 63.32 | -5.64 | 9.78 |
| × | ✓ | ✓ | 67.32 | 62.41 | -15.23 | 13.32 | 62.17 | 58.14 | -16.67 | 18.68 |
| ✓ | × | ✓ | 76.04 | 73.44 | -3.95 | 2.23 | 68.52 | 64.12 | -8.30 | 6.73 |
| ✓ | ✓ | × | 80.35 | 79.68 | -1.21 | 0.80 | 69.78 | 63.48 | -2.13 | 3.46 |
| ✓ | ✓ | ✓ | 82.13 | 78.42 | -0.1077 | 0.17 | 72.79 | 68.93 | -0.1805 | 0.7923 |

The full model, which integrates retrieval, LoRA composition, and prompts, achieves the highest accuracy and the lowest forgetting. These results suggest that retrieval and adapter mixing are the dominant contributors, while prompts play a complementary role by refining early-stage protein representations.

We first analyze the behavior of the prototype-based retrieval module on the BioSNAP dataset. Figure 4(a) shows the domain-level retrieval confusion matrix, which exhibits a strong diagonal pattern: proteins most frequently retrieve prototypes originating from their own domains. Figures 4(b-c) further illustrate that protein embeddings cluster tightly around their associated prototypes in the projected space, confirming that the prototypes faithfully summarize domain structure.

**Fig. 4:**
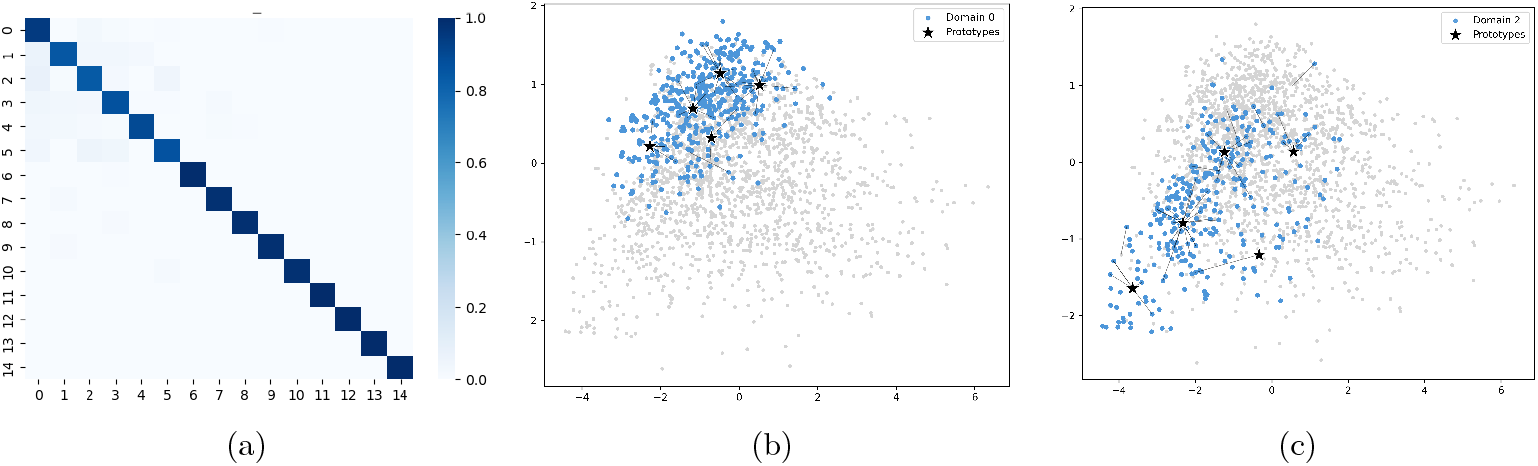
Prototype retrieval visualization on BIOSNAP. (a) Cross-domain retrieval confusion matrix, where each entry indicates the proportion of proteins in one domain whose top-1 retrieved prototype belongs to another domain. (b, c) Two-dimensional projections of protein embeddings and prototypes for two representative domains. Blue points represent proteins from the target domain, gray points represent proteins from other domains, and black stars denote stored prototypes. Arrows indicate the retrieved nearest prototypes.

To evaluate the reliability of the matching process, we compare several similarity measures and find that Manhattan distance produces the highest top-1 retrieval accuracy, reaching 93.12%. Additional results are provided in Appendix E.

We also assess computational efficiency. The prototype memory stores five 1280-dimensional float32 vectors for each domain, which corresponds to roughly 25.6 KB of storage and stays below 1 MB even with 25 domains. Retrieval is highly efficient as well; using FAISS, each query takes less than 2 ms and accounts for under 5 percent of total inference time.

### 3.4 Interpretability Analysis

To assess the interpretability and biological alignment of PRADA-DTI, we visualize residue-level attribution maps for proteins from four different domains. Figure 5 displays attribution intensities on the solvent-accessible surface, rendered on a blue-white-red scale (from low to high importance). Across all domains, regions with strong attribution consistently align with ligand-binding pockets, demonstrating that the model reliably identifies functional sites.

**Fig. 5:**
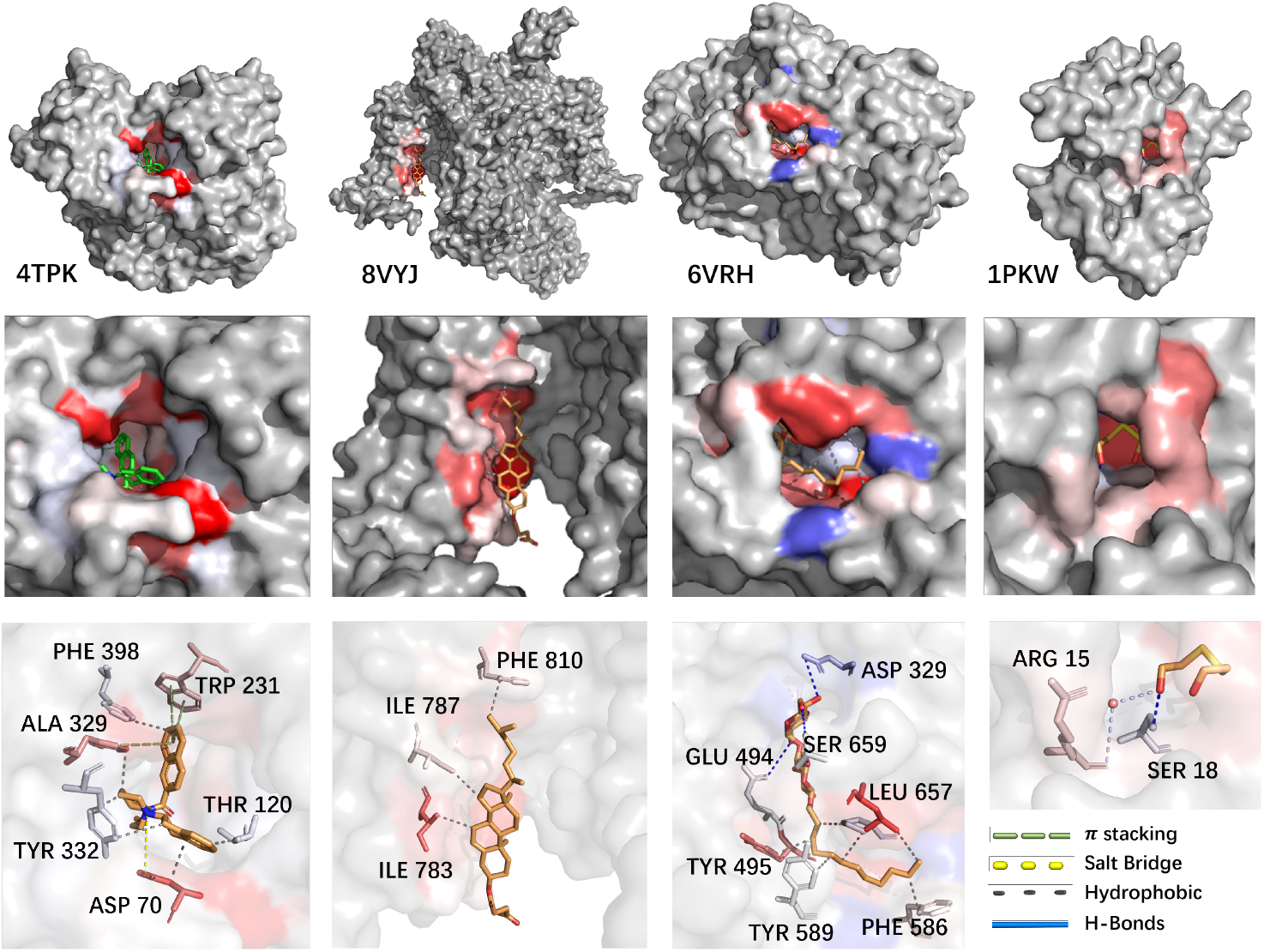
Residue-level attribution maps for proteins from four different domains. The first row shows the overall protein surfaces; the second row highlights the local ligand-binding pockets with attribution intensity rendered on a blue–white–red scale (from low to high importance); the third row presents detailed protein–ligand interactions, including hydrogen bonds, hydrophobic contacts, aromatic stacking, and salt bridges. PRADA-DTI consistently focuses on the true binding pockets across all domains.

To further assess biological relevance, we annotate binding residues and interaction types using the PLIP interaction profiler [25]. The structural views in Figure 5 show that PRADA-DTI highlights residues involved in a wide range of protein-ligand interactions, including hydrogen bonds (SER, GLU, ASP), hydrophobic contacts (PHE, TYR, ILE, LEU), aromatic stacking (TRP, ALA), and salt bridges (ASP). The attribution maps closely match experimentally validated binding sites, confirming that the model’s focus corresponds to meaningful biochemical structure rather than nonspecific surface regions.

These results demonstrate that PRADA-DTI produces clear and biologically grounded attribution signals under continual learning, and generalizes pocket-recognition behavior across diverse protein domains. This highlights the method’s strong interpretability and its potential utility for explainable drug-target interaction prediction.

## 4 Conclusion

To address the challenge of catastrophic forgetting and domain shifts in drug-target interaction prediction, we propose PRADA-DTI, a retrieval-augmented and domain-adaptive framework for continual learning. By integrating prototype-guided retrieval, prompt-based domain conditioning, and dynamically composed LoRA adapters, PRADA-DTI enables privacy-preserving, parameter-efficient, and task-ID-free adaptation across protein domains. Experiments on BIOSNAP and BindingDB demonstrate that our method significantly improves predictive accuracy and forgetting resistance while updating less than 5% of backbone parameters, establishing a new paradigm for continual molecular modeling in computational drug discovery.

## Supporting information

all supplemental materials

## Code Availability

All source code used in this study, along with documentation and a minimal test dataset, is publicly available at:https://github.com/LastWhisper114514/PRADA-DTI

## Acknowledgments

This work is supported in part by the National Natural Science Foundation of China under grants W2411054, U21A20521, and 62271178; the Postgraduate Research and Practice Innovation Program of Jiangsu Province (KYCX23_2524); the National Foreign Expert Project of China under Grant G2023144009L; the Zhejiang Provincial Natural Science Foundation of China (LR23F010002); the Wuxi Health Commission Precision Medicine Project (J202106); the Jiangsu Provincial Six Talent Peaks Project (YY-124); and the Major Projects of Wuxi Health Commission (Z202324).

## Disclosure of Interests

The authors declare that they have no competing interests related to the content of this article.

## Notes

### Competing Interest Statement

The authors have declared no competing interest.

## Bibliography

[1] Ozturk, H., Ozkirimli, E., Ozgur, A.: DeepDTA: deep drug-target binding affinity prediction. Bioinformatics 34(17), i821–i829 (2018)

[2] Huang, K., Fu, T., Xiao, C., Glass, L. M., Sun, J., Zitnik, M., Gomes, C. P.: MolTrans: Molecular interaction transformer for drug-target interaction prediction. Bioinformatics 37(6), 830–836 (2021)

[3] Bleakley, K., Yamanishi, Y.: Supervised prediction of drug-target interactions using bipartite local models. Bioinformatics 25(18), 2397–2403 (2009)

[4] Yamanishi, Y., Araki, M., Gutteridge, A., Honda, W., Kanehisa, M.: Prediction of drug-target interaction networks from the integration of chemical and genomic spaces. Bioinformatics 24(13), i232–i240 (2008)

[5] H. Öztürk, A. Özgür, and E. Ozkirimli. DeepDTA: deep drug–target binding afinity prediction. Bioinformatics, 34(17):i821–i829, 2018.

[6] K. Huang, C. Xiao, L. M. Glass, and J. Sun. MolTrans: Molecular interaction transformer for drug–target interaction prediction. Bioinformatics, 37(6):830–836, 2021.

[7] Nguyen, T., Le, H., Quinn, T. P., Nguyen, T., Le, T. D., Venkatesh, S.: GraphDTA: Predicting drug-target binding affinity with graph neural networks. Bioinformatics 37(8), 1140–1147 (2021)

[8] Chen, L., Tan, X., Wang, D., Zhong, F., Liu, Z., Zeng, X.: TransformerCPI: improving compound-protein interaction prediction by sequence-based deep learning. Briefings in Bioinformatics 22(6), bbab158 (2021)

[9] Wang, Y., You, R., Chen, X., Liu, F., Yan, Z.: DeepPurpose: A deep learning library for drug-target interaction prediction. Bioinformatics 38(11), 3258–3260 (2022)

[10] Liu, R., Liu, Z., Zhao, Y., et al.: DTIC: A benchmark for deep learning-based drug-target interaction prediction under class imbalance and cold-start scenarios. Briefings in Bioinformatics, bbac540 (2023)

[11] Huang, K., Fu, T., Gao, W., et al.: Therapeutics Data Commons: Machine learning datasets and tasks for drug discovery and development. arXiv preprint 2102.09548 (2021)

[12] Hsu, Y. C., Liu, Y. C., Ramasamy, A., Kira, Z.: Re-evaluating continual learning scenarios: A categorization and case for strong baselines. In: NeurIPS (2018)

[13] van de Ven, G. M., Tolias, A. S.: Three scenarios for continual learning. arXiv preprint 1904.07734 (2019)

[14] Kirkpatrick, J., Pascanu, R., Rabinowitz, N., et al.: Overcoming catastrophic forgetting in neural networks. PNAS 114(13), 3521–3526 (2017)

[15] Lopez-Paz, D., Ranzato, M.: Gradient episodic memory for continual learning. In: NeurIPS (2017)

[16] Rusu, A. A., Rabinowitz, N. C., Desjardins, G., et al.: Progressive neural networks. arXiv preprint 1606.04671 (2016)

[17] Li, J., Wang, X., Xie, X.: MoleCL: Continual learning for molecular property prediction. arXiv preprint 2206.05048 (2022)

[18] Wang, Y., Jin, X., Hou, Y., et al.: Graph contrastive continual learning for molecular property prediction. In: ICLR (2023)

[19] Hu, E. J., Shen, Y., Wallis, P., et al.: LoRA: Low-rank adaptation of large language models. In: ICLR (2022)

[20] Lester, B., Al-Rfou, R., Constant, N.: The power of scale for parameter-efficient prompt tuning. In: EMNLP (2021)

[21] Jia, H., Lin, X., Chen, X., et al.: Visual prompt tuning. In: ECCV (2022)

[22] Lewis, P., Perez, E., Piktus, A., et al.: Retrieval-augmented generation for knowledge-intensive NLP tasks. In: NeurIPS (2020)

[23] Z. Lin, H. Akin, R. Rao, B. Hie, Z. Zhu, W. Lu, N. Smetanin, R. Verkuil, O. Kabeli, Y. Shmueli, A. D. S. Costa, M. Fazel-Zarandi, T. Sercu, S. Candido, and A. Rives, “Evolutionary-scale prediction of atomic-level protein structure with a language model,” Science, vol. 379, no. 6637, pp. 1123– 1130, 2023. DOI: 10.1126/science.ade2574.

[24] S. Chithrananda, G. Grand, and B. Ramsundar, “ChemBERTa: Large-scale self-supervised pretraining for molecular property prediction,” arXiv preprint 2010.09885, 2020.

[25] M. F. Adasme, K. L. Linnemann, S. N. Bolz, F. Kaiser, S. Salentin, V. J. Haupt, and M. Schroeder, “PLIP 2021: expanding the scope of the protein-ligand interaction profiler to DNA and RNA,” Nucleic Acids Research, vol. 49, no. W1, pp. W530–W534, 2021. DOI: 10.1093/nar/gkab294.

[26] Kirkpatrick, J., Pascanu, R., Rabinowitz, N., Veness, J., Desjardins, G., Rusu, A. A., Milan, K., Quan, J., Ramalho, T., Grabska-Barwiska, A., Hassabis, D., Clopath, C., Kumaran, D., and Hadsell, R. Overcoming catastrophic forgetting in neural networks. Proceedings of the National Academy of Sciences (PNAS), 114(13):3521–3526, 2017.

[27] Li, Z. and Hoiem, D. Learning without forgetting. In European Conference on Computer Vision (ECCV), pp. 614–629. Springer, 2016.

[28] B. Lester, R. Al-Rfou, and N. Constant, “The Power of Scale: Parameter-Efficient Adaptation for Pretrained Language Models,” in Proceedings of the 2021 Conference on Empirical Methods in Natural Language Processing (EMNLP), pp. 3045–3059, 2021. DOI: 10.18653/v1/2021.emnlp-main.243.

[29] Xie, Y., Guo, Z., Zhang, L., et al. Task-specific Parameter Partitioning for Lifelong Molecular Property Prediction. Advances in Neural Information Processing Systems (NeurIPS), 2022.

[30] X. Zhang, D. Song, and D. Tao, “Hierarchical Prototype Networks for Continual Graph Representation Learning,” IEEE Transactions on Pattern Analysis and Machine Intelligence, vol. 45, no. 4, pp. 4622–4636, Apr. 2023. DOI: 10.1109/TPAMI.2022.3186909.

[31] J. Cai, X. Wang, C. Guan, Y. Tang, J. Xu, B. Zhong, and W. Zhu, “Multimodal Continual Graph Learning with Neural Architecture Search,” in Proceedings of the ACM Web Conference 2022 (WWW ‘22), Virtual Event, Lyon, France, pp. 1292–1300, 2022. DOI: 10.1145/3485447.3512176.

