## Supplementary material for "PRADA-DTI: A Prototype-Retrieval Augmented Domain-Adaptation Framework for Drug-Target Interaction Prediction": all supplemental materials

### Appendix A.Experiment Parameter Details

We performed a grid search over the Leiden clustering hyperparameters, including the *resolution* of the Constant Potts Model (CPM) and the number of neighbors (*kNN*) used to construct the graph. For each parameter combination, we evaluated clustering quality using the following metrics:

- **silhouette**: higher values indicate better cluster separation.
- **davies_bouldin**: a lower value indicates more compact and well-separated clusters.
- **modularity**: measures the clarity of graph communities; higher is better.
- **score**: our composite metric defined as

$$\text{score} = \text{Silhouette} - 0.2 \times \text{Davies–Bouldin} + 2 \times \text{Modularity}.$$

- **n_clusters**: number of clusters produced by the Leiden algorithm.

The complete results of the grid search are shown in Table below:

| resolution | knn | silhouette | davies_bouldin | modularity | score | n_clusters |
|---|---|---|---|---|---|---|
| 0.001 | 10 | -0.06 | 2.40 | 0.26 | -0.03 | 8 |
| 0.001 | 20 | 0.08 | 2.22 | 0.09 | -0.18 | 2 |
| 0.001 | 30 | -1.00 | 999.00 | 0.00 | -200.80 | 1 |
| 0.001 | 40 | -1.00 | 999.00 | 0.00 | -200.80 | 1 |
| 0.005 | 10 | 0.02 | 2.61 | 0.71 | 0.92 | 15 |
| 0.005 | 20 | -0.03 | 2.70 | 0.50 | 0.43 | 6 |
| 0.005 | 30 | 0.03 | 2.26 | 0.11 | -0.20 | 3 |
| 0.005 | 40 | 0.08 | 2.22 | 0.08 | -0.21 | 2 |
| 0.01 | 10 | 0.02 | 2.73 | 0.72 | 0.92 | 24 |
| 0.01 | 20 | 0.04 | 2.59 | 0.64 | 0.80 | 11 |
| 0.01 | 30 | 0.05 | 2.92 | 0.53 | 0.53 | 5 |
| 0.01 | 40 | 0.07 | 3.14 | 0.41 | 0.27 | 3 |
| 0.1 | 10 | 0.08 | 2.10 | 0.60 | 0.87 | 137 |
| 0.1 | 20 | 0.07 | 2.21 | 0.54 | 0.70 | 81 |
| 0.1 | 30 | 0.06 | 2.49 | 0.50 | 0.56 | 51 |
| 0.1 | 40 | 0.03 | 2.22 | 0.50 | 0.58 | 45 |
| 0.2 | 10 | 0.11 | 1.83 | 0.53 | 0.81 | 240 |
| 0.2 | 20 | 0.09 | 1.88 | 0.47 | 0.66 | 156 |
| 0.2 | 30 | 0.07 | 1.95 | 0.44 | 0.55 | 123 |
| 0.2 | 40 | 0.05 | 1.99 | 0.42 | 0.49 | 97 |
| 0.5 | 10 | 0.15 | 1.39 | 0.44 | 0.74 | 466 |
| 0.5 | 20 | 0.12 | 1.52 | 0.37 | 0.55 | 335 |
| 0.5 | 30 | 0.11 | 1.61 | 0.33 | 0.44 | 273 |
| 0.5 | 40 | 0.09 | 1.65 | 0.30 | 0.36 | 242 |
| 1 | 10 | 0.15 | 0.94 | 0.33 | 0.63 | 821 |
| 1 | 20 | 0.14 | 1.06 | 0.28 | 0.48 | 648 |
| 1 | 30 | 0.13 | 1.16 | 0.24 | 0.37 | 554 |
| 1 | 40 | 0.12 | 1.23 | 0.21 | 0.29 | 494 |
| 1.5 | 10 | 0.15 | 0.89 | 0.26 | 0.49 | 963 |
| 1.5 | 20 | 0.14 | 1.01 | 0.21 | 0.36 | 785 |
| 1.5 | 30 | 0.13 | 1.08 | 0.18 | 0.27 | 702 |
| 1.5 | 40 | 0.12 | 1.15 | 0.15 | 0.20 | 639 |

Table 3: Grid search results for Leiden clustering. Each row corresponds to one (resolution, kNN) setting and the associated clustering evaluation metrics.

Based on the composite score, the best-performing configuration is:

$$\textbf{resolution} = 0.005, \qquad \textbf{kNN} = 10, \qquad \textbf{score} = 0.924.$$

This configuration is therefore adopted in our clustering pipeline, yielding **15 clusters on Biosnap** and **25 clusters on BindingDB**.

To determine the optimal configuration for prototype-based retrieval, we conducted a grid search over the number of stored prototypes (n_proto) and the number of retrieved prototypes $k$. Table tab:placeholder presents the resulting accuracy and average forgetting across various settings.

| n_proto | k | ACC | AFGT | n_proto | k | ACC | AFGT |
|---|---|---|---|---|---|---|---|
| 1 | 1 | 75.52 | 10.46 | 7 | 5 | 78.75 | 3.48 |
| 3 | 1 | 76.78 | 1.25 | 7 | 7 | 76.79 | 7.45 |
| 3 | 3 | 77.15 | 0.72 | 9 | 1 | 79.13 | 1.25 |
| 5 | 1 | 77.43 | 1.59 | 9 | 3 | 81.42 | 0.85 |
| 5 | 3 | **82.13** | **0.17** | 9 | 5 | 81.42 | 0.24 |
| 5 | 5 | 78.06 | 5.88 | 9 | 7 | 76.43 | 8.67 |
| 7 | 1 | 78.82 | 2.30 | 9 | 9 | 70.48 | 13.33 |
| 7 | 3 | 81.25 | 1.35 | - | - | - | - |

Table 4: Evaluation of prototype retrieval configurations. Accuracy (ACC) and average forgetting (AFGT) are reported for varying numbers of stored prototypes $n\_proto$ and retrieved neighbors $k$. The best-performing setting is highlighted **in bold**.

We observe that increasing n_proto generally improves performance up to a certain point, with diminishing returns or degradation beyond that. Notably, the configuration with n_proto=5 and k=3 achieves the highest accuracy of 82.13% and the lowest AFGT at 0.17, indicating both strong performance and stability during continual learning. Larger k values sometimes lead to higher AFGT, suggesting potential over-reliance on distant prototypes. Based on these results, we adopt n_proto = 5 and k = 3 in all subsequent experiments.

### Appendix B. Parameter Sensitivity Analysis

To examine how model parameters evolve during continual domain adaptation, we visualize the distribution of parameter changes between the first and the last domains. Figure 6 shows the histogram of Frobenius-norm differences across all trainable parameters.

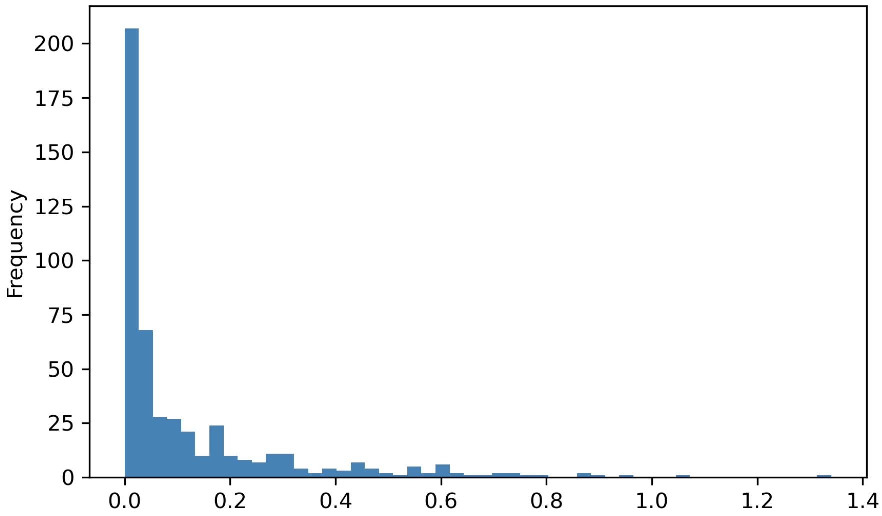

Fig. 6: **Distribution of parameter changes between the first and last domains.** The x-axis represents the magnitude of parameter updates (Frobenius norm), while the y-axis indicates the number of parameters within each interval. Most parameters exhibit negligible changes (near zero), whereas only a small subset shows large updates. This long-tailed distribution implies that adaptation is concentrated in a few sensitive layers, supporting the feasibility of inserting LoRA modules into limited key layers rather than uniformly across the entire network.

These results demonstrate that our method achieves domain adaptation mainly through localized parameter adjustments. Therefore, the model can maintain high efficiency by applying LoRA-based fine-tuning only to layers that are empirically identified as adaptation-sensitive, without sacrificing performance or requiring full-parameter updates.

### Appendix C. Visualization of Parameter Change Distribution

To complement the statistical view in Figure 6, we visualize the spatial distribution of parameter changes across all layers. Figure 7 presents a one-dimensional heatmap illustrating the Frobenius-norm differences of each parameter between the first and last domains.

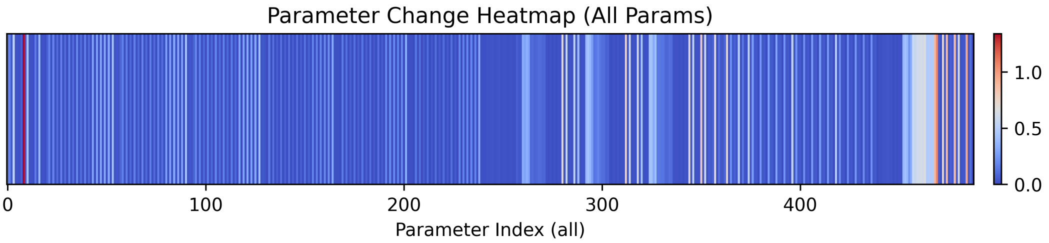


Fig. 7: **Parameter change heatmap across all layers.** Each vertical stripe represents a parameter (or parameter group) arranged in model order. The color intensity encodes the magnitude of parameter updates, where blue denotes negligible change and red indicates large variation. The sparsity of red regions suggests that only a few parameter blocks undergo substantial modification during adaptation, further confirming that LoRA modules can be inserted selectively into these sensitive layers rather than uniformly throughout the network.

Together with the histogram in Figure 6, this visualization highlights that domain adaptation in our model is achieved through localized and sparse parameter shifts, demonstrating the efficiency and feasibility of partial LoRA integration.

### Appendix D. Cross-Domain Protein Attribution Visualization

To further investigate how domain-specific prompts influence residue-level importance, we visualize the attribution maps of the same protein across all domains. Figure 8 shows that attention patterns vary considerably between domains, indicating that the model indeed learns domain-conditioned protein representations. Such variability justifies the need for domain-specific prompt tuning in our framework.

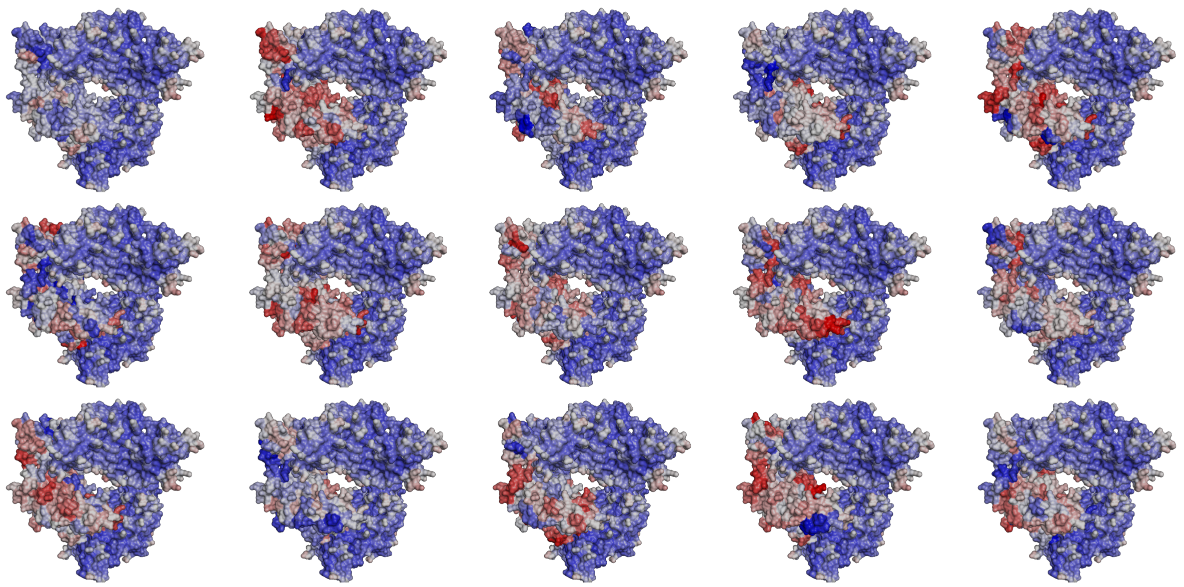

Fig. 8: **Residue-level attribution maps across 15 domains for the same protein.** Colors denote attention intensity from low (blue) to high (red). The large cross-domain variation highlights how domain-conditioned prompts reshape protein representations during domain adaptation.

### Appendix E. Comparison of Similarity Measures for Prototype Retrieval

To evaluate the robustness of prototype retrieval, we tested multiple distance and similarity metrics, including cosine similarity, Manhattan distance, Euclidean distance, correlation distance, and several kernel-based similarities. The Top-1 retrieval accuracies under different prototype counts ($n_{\text{proto}} \in \{1, 3, 5, 7, 9\}$) are summarized in Figure 9.

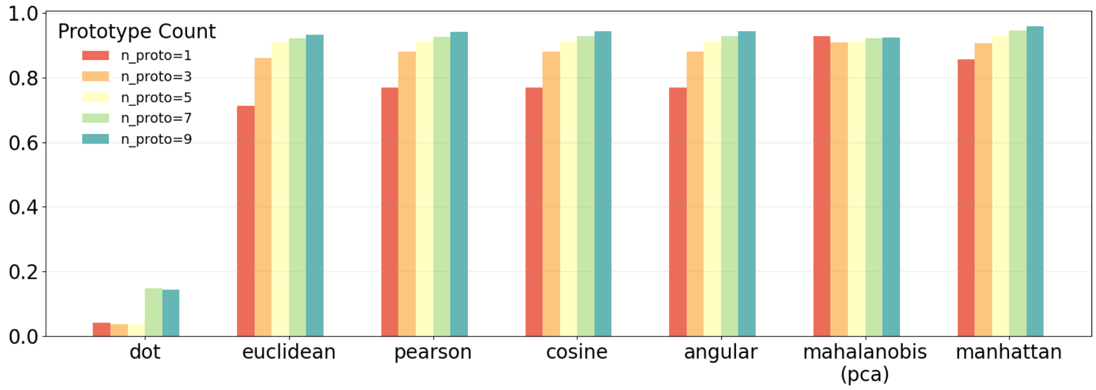


Fig. 9: **Prototype retrieval performance under different similarity metrics.** The x-axis lists similarity or distance measures sorted by mean Top-1 accuracy. Each color denotes a different number of prototypes per domain.

These results confirm that cosine similarity is a stable and effective metric for prototype matching, and justify its selection as the default retrieval formulation in our framework.
